# EpiVar Browser: advanced exploration of epigenomics data under controlled access

**DOI:** 10.1101/2023.08.03.551309

**Authors:** David R Lougheed, Hanshi Liu, Katherine A Aracena, Romain Grégoire, Alain Pacis, Tomi Pastinen, Luis B Barreiro, Yann Joly, David Bujold, Guillaume Bourque

## Abstract

**Motivation:** Human epigenomic data has been generated by large consortia for thousands of cell types to be used as a reference map of normal and disease chromatin states. Since epigenetic data contains potentially identifiable information, similarly to genetic data, most raw files generated by these consortia are stored in controlled-access databases. It is important to protect identifiable information, but this should not hinder secure sharing of these valuable datasets.

**Results:** Guided by the *Framework for responsible sharing of genomic and health-related data* from the Global Alliance for Genomics and Health (GA4GH), we have developed a tool to facilitate the exploration of epigenomics datasets’ aggregate results, while filtering out identifiable information. Specifically, the EpiVar Browser allows a user to navigate an epigenetic dataset from a cohort of individuals and enables direct exploration of genotype-chromatin phenotype relationships. Because the information about individual genotypes is not accessible and aggregated in the output that is made available, no identifiable data is released, yet the interface allows for dynamic genotype - epigenome interrogation. This approach has the potential to accelerate analyses that would otherwise require a lengthy multi-step approval process and provides a generalisable strategy to facilitate responsible access to sensitive epigenomics data.

**Availability and implementation:** Online portal instance: https://computationalgenomics.ca/tools/epivar

Source code: https://github.com/c3g/epivar-browser

## 1 Introduction

In recent years, multiple international consortia have been coordinating the mapping of the epigenetic landscape across human tissues and individuals. They include the Encyclopedia of DNA Elements consortium (ENCODE) [ENCODE], the NIH Roadmap [Roadmap] and the International Human Epigenome Consortium (IHEC) [IHEC], which together have profiled the epigenetic features of over 3000 biosamples, in over 8000 epigenomic experiments. The hope is that these detailed epigenetic maps will lead to a better understanding of the human genome, including a better annotation of non-coding regulatory elements. The Genotype-Tissue Expression (GTEx) project goes a step further and is characterising the effects of genetic variation on gene expression and chromatin [GTEx]. Through the identification of quantitative trait loci (QTLs) with either expression (eQTL) or chromatin (cQTL), the aim is to improve our understanding of the molecular mechanisms of genetic risk for complex traits and diseases. Notably, in line with open science principles and to facilitate further analyses and discoveries, the cited initiatives all provide access to various summary data on their portal. However, one of the main issues for researchers interested in these resources is that developing new visualisations or performing novel analyses on them can be challenging [sci-data]. That’s because the underlying raw data is hosted in registered or controlled-access databases, to protect potentially re-identifiable genetic data. In this context, we need better mechanisms to facilitate epigenomic data discovery and analysis, while addressing the ethical and privacy aspects associated with data sharing [Sharing].

Focusing mainly on genetic data, the Global Alliance for Genomics and Health (GA4GH) has been working on secure methods federating sensitive genomic data to facilitate data discovery. This includes the development of the Beacon v2 API specification [beacon v2], defining a programming interface that enables third party tools and portals to discover and query the clinical and genomics content of project-specific databases. GA4GH has now also started to add API specifications for other -omics data, such as from RNA-seq experiments [rnaget]. Beyond supporting epigenomics data discovery, we wanted to enable data integration in a manner consistent with the GA4GH *Framework for responsible sharing of genomic and health-related data* [framework]. Specifically, we wanted to facilitate genotype-chromatin phenotype relationships such as the ones identified by GTEx or IHEC [flu-paper]. Once a QTL is identified, we would facilitate the exploration of that locus and look at the underlying epigenomics data supporting an association. Currently, this can only be accomplished by a lengthy multistep process involving a data access-request, approval, download and an ad hoc analysis to group epigenomics tracks based on genotype. With the EpiVar Browser, we propose a seamless solution that combines the epigenomics data from a cohort based on genotype and allows direct visualisation of the tracks. Using the D-PATH policy assessment tool [D-PATH], we ensured no identifiable information is released, meaning the data can be provided without a data access request. We still require a login to ensure that data use conditions are acknowledged and respected.

## 2 Results

We have implemented the EpiVar Browser, a portal that stores access-controlled genetic and epigenetic datasets and enables researchers to explore the interaction between genomic variants and epigenetic features, such as histone tail modifications, chromatin accessibility, and the transcriptome. As a proof of concept, we used data with 510 epigenomics experiments from six different assays and 35 individuals obtained from a study exploring the response to influenza infection [flu-paper]. This dataset contains over 5 million single nucleotide polymorphisms (SNPs) associated with ∼376 000 genomic features for a total of around 29 million associations at p < 0.05. A web interface allows users to provide a variant of interest, using its dbSNP accession ID (“rs” number) [dbsnp], and suggests genomic features with the most significant associations (Figure S1). Alternatively, a user can provide the symbol for a gene of interest, after which they can choose a variant with relevant detected associations. The tool generates two boxplots per epigenetic assay available, stratified by genotype for that variant, and by flu infection status (Figure 1A). The list of conditions (here, flu/non-infected) and population groups (here, ethnicity) can be configured on the server-side by portal administrators. Users can also visualise results in the UCSC Genome Browser [UCSC], with tracks displaying the average of the experiment signal per genotype (Figure 1B). These tracks are calculated upon request for a 200 kilobases (kb) genomic window centred on the feature of interest.

**Figure 1.**
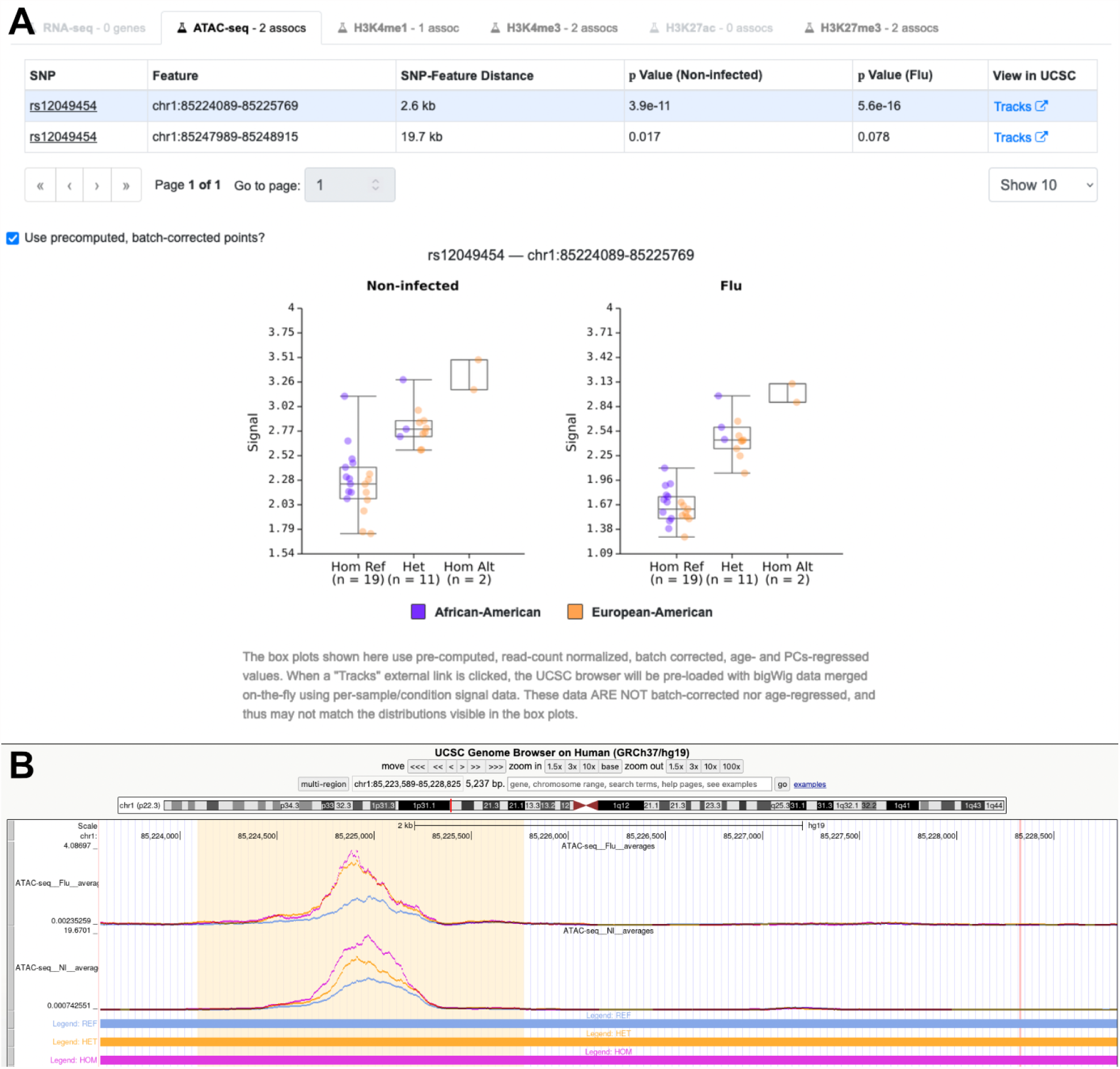
The EpiVar browser generates summary data box plots on-the-fly using sample bigWig files for a selected assay and genotype information on a secure web server (A). These plots, in this case, reveal correlations between the rs12049454 SNP genotype and chromatin accessibility in both non-infected and flu-infected cells. The merged signal for the feature (yellow highlighted region), stratified by genotype (SNP locus highlighted in red), can be visualised in the UCSC Genome Browser (B).

## 3 Implementation and compliance

The code for the EpiVar Browser is open source, and released under the LGPL version 3.0 licence [github]. The box plot charts are either created from precomputed point matrices or dynamically generated for the user-selected feature using the UCSC *bigWigSummary* tool [bigWig]. In either case, box plots are derived from locally-stored signal and genotype data. Genotypes for all participants are stored in a GEMINI database [GEMINI].

Signal tracks for the all epigenetic experiments (one per assay/participant) are stored on the EpiVar server in the bigWig format. As much as possible, all data processing is done close to the source (i.e., server-side) – both on-the-fly data processing [bigWig] and plot generation. Association significance for peak-SNP pairs must be computed ahead of time and imported into the portal. Precomputed signal values (to incorporate, e.g., batch correction) for feature-sample pairs can optionally be provided as well.

Browsing of tracks on the UCSC Genome Browser is made possible by producing a Track Hub with dynamically generated bigWig files for the genomic region of interest, per assay and per average signal for a genotype group. The original per-sample bigWig files were normalised so that each value represents the read count per base pair per 10 million reads. A Node.js application and a signal annotation file-processing service efficiently slices these bigWig files and merges them on demand, using a modified version of the *bigWigMerge* utility from UCSC. Genome browser tracks were created with the HOMER *makeUCSCfile* command and *bedGraphToBigWig* utility from UCSC. These tracks are created on the fly by averaging bigWig regions of samples sharing an experimental treatment and genotype. The track’s curve denotes the distribution of the average RPM values. A second track hub, pre-configured by portal administrators, shows signals of various epigenetic marks coloured by condition.

During the implementation of the portal, we put particular attention into protecting the potentially privacy-compromising aspects of the data. This is why some design decisions were taken as to how data is displayed (e.g., box plots generated at the server-side, rather than presenting an API which may enumerate samples in a specific order). The EpiVar Browser also doesn’t reveal individuals’ signal level/genotype pair with identifiers, so that the genotype data cannot be collected across multiple SNPs. All of this is to prevent the re-identification of individuals using genetic information. Finally, we used the D-PATH privacy assessment tool to identify the applicable legal and ethical requirements regarding the sharing of epigenetic data and accompanying metadata [D-PATH].

The raw data for genomic and epigenetic analyses is deposited at the European Genome-Phenome Archive (EGA). Researchers with an interest to get more information, such as participant-level granularity of the data, can apply for access to the relevant Data Access Committee.

## 4 Discussion

With the EpiVar Browser, potentially privacy-compromising data such as per-individual genotypes, usually stored under controlled access, are never directly accessible to the portal users; instead, they are processed to generate summary data on demand. Variant information is instead made available by genotype groups. Even though we put our best efforts in protecting participant-level data to minimise the risk of re-identification, registered access is required in order to make users agree to a terms of use which forbids participants re-identification attempts.

The code for the EpiVar Browser could be used to implement other portals to provide access to various epigenomic datasets and allow the efficient exploration of other genotype-phenotype associations. Finally, what we presented here also represents a strategy that could be used more generally to accelerate access to sensitive epigenomics and other datasets.

## Acknowledgements

We would like to thank Calcul Québec and the Digital Research Alliance of Canada for access to expertise along with compute and storage resources.

## Funding

This work was supported by a Canada Institute of Health Research (CIHR) program grant (CEE-151618) for the McGill Epigenomics Mapping Center, which is part of the Canadian Epigenetics, Environment and Health Research Consortium (CEEHRC) Network. The work on EpiVar was also supported by a Genome Canada grant called EpiShare (Genome Canada - 15502) and a Technology Platform grant for the Canadian Center for Computational Genomics (C3G). This project was also supported by National Institute of Health Research grants R01-GM134376 and P30-DK042086 to L.B.B. GB is supported by a Canada Research Chair Tier 1 award, a Fonds de recherche du Québec - Santé (FRQ-S) Distinguished Research Scholar award, and by the World Premier International Research Center Initiative (WPI), MEXT, Japan. K.A.A. is supported by a grant to University of Chicago from the Howard Hughes Medical Institute through the James H. Gilliam Fellowships for Advanced Study program.

## Data availability

All the data used for the EpiVar Browser is available under controlled access at the European Genome Phenome Archive, with the dataset IDs EGAD00001008422 and EGAD00001008359.

## Supplementary figures

**Figure S1:**
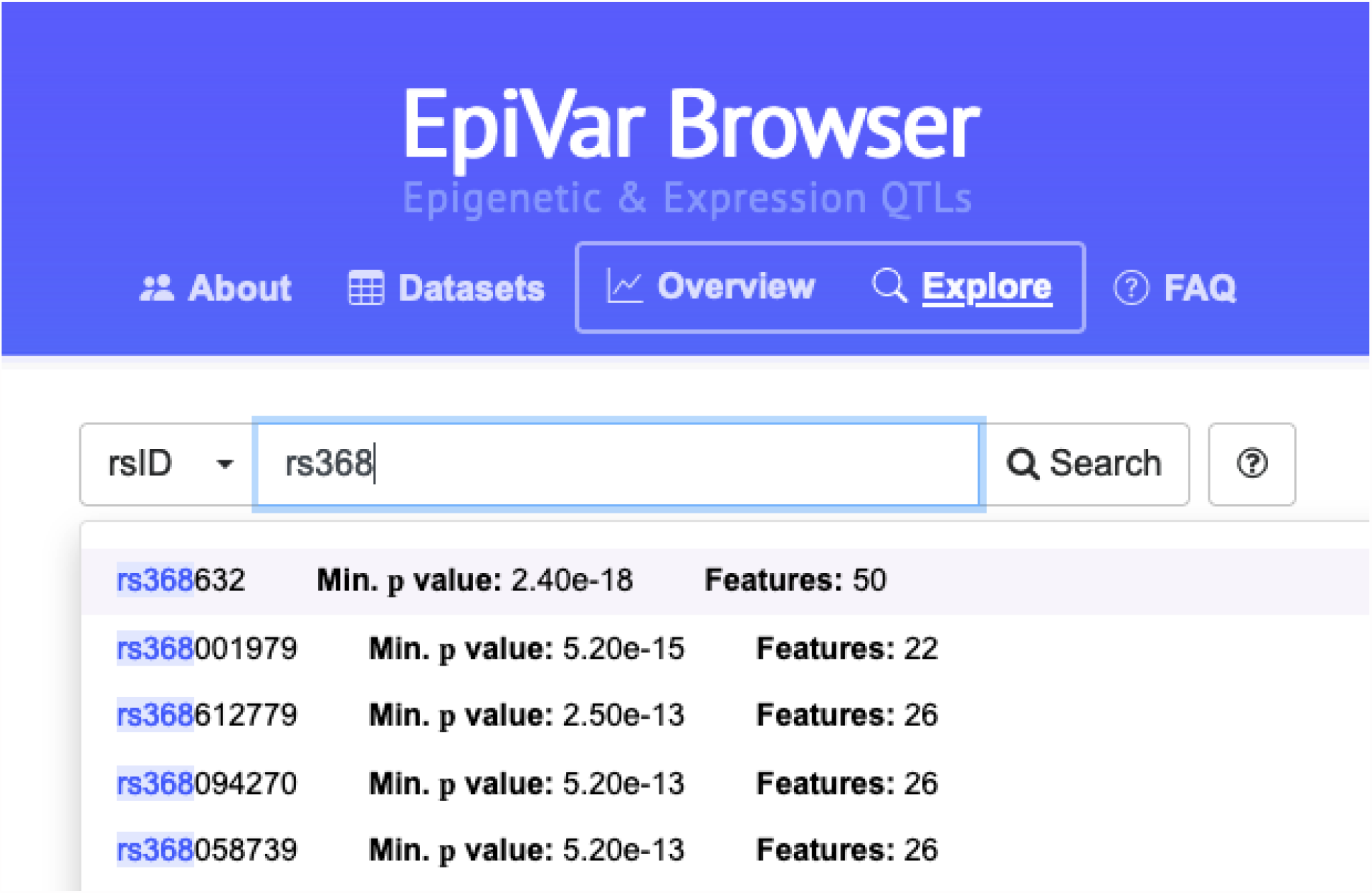
The EpiVar Browser allows SNP searching via rsID. Up to 50 matching results are shown; SNPs with the strongest available feature associations are shown first. Selecting a SNP will take a user to a page with the strongest peak, and the assay it came from, pre-selected.

## Notes

https://computationalgenomics.ca/tools/epivar

https://github.com/c3g/epivar-browser

### Competing Interest Statement

The authors have declared no competing interest.

